# *Arginine Kinase 1* regulates energy homeostasis in *Drosophila* muscle development

**DOI:** 10.64898/2026.01.30.702107

**Authors:** Maria Paula Zappia, Anton Westacott, Hannah Cooke, Rhianna Geary, Libby Travers, Lucia de Castro, Oliver Carty, Maxim V Frolov

## Abstract

In *Drosophila*, Arginine kinase 1 (Argk1) is involved in maintaining ATP homeostasis during bursts of activity in tissues with high and variable rates of energy turnover such as muscle. However, its role beyond stress conditions is less understood. Here, we show that *Argk1* maintains energy homeostasis during flight muscle development and is required for animal viability and proper muscle function. The knockdown of Argk1 causes defects in both early and late stages of myogenesis. In the proliferating myoblasts associated with the wing disc, Argk1 depletion results in a reduction in cell size without changes in cell cycle progression. Single cell RNA-sequencing revealed that the overall composition of differentiating and undifferentiating myoblasts is not altered. Nonetheless, Argk1 knockdown causes broad alterations in the expression of genes involved in various metabolic pathways. This correlates with low levels in both ATP content and NAD+/NADH ratio. Later in muscle development, Argk1-depleted muscles completely lack spontaneous muscle contractions that are essential in myofibrillogenesis. Accordingly, Argk1 knockdown results in severe defects in sarcomere structure, while the mitochondrial network is highly fragmented. Furthermore, muscle growth is severely reduced. Thus, our data reveal an essential role for Argk1 in maintaining energy homeostasis throughout muscle development, which is required to meet the demand to support myofibrillogenesis, muscle growth and proper muscle function.

## Introduction

The *Drosophila* model organism has been extensively used to elucidate the fundamental genetic mechanisms and biological processes underlying muscle development. Somatic fly muscles closely resemble vertebrate skeletal muscles, as both share highly conserved molecular network and the signaling pathways (*1*, *2*). The development of the skeletal muscles follows a precise stepwise process to build fully functional muscles that can support all kinds of locomotion, including flying, walking, jumping and crawling (*3*). The *Drosophila* adult skeletal muscles are formed during the second wave of myogenesis later in development (*4*). Briefly, the adult muscle precursor (AMP) cells or myoblasts that will give rise to the flight muscles are specified by asymmetric division of the muscle progenitors during embryonic development and remain quiescent until larval stages when they are activated and rapidly proliferate to form a layer of adepithelial cells tightly associated with the wing imaginal discs. Then, the reduction in the expression of the transcription factor Twist and decrease of Notch signaling results in the activation of the myogenic differentiation program (*5*, *6*) leading to the myoblast fusion, followed by muscle growth, myofibrillogenesis and sarcomere assembly during pupal development. There are two types of flight muscles, the direct flight muscles (DFM) and the indirect flight muscles (IMF). These two flight muscles show distinct physiology, structure and function. The IFM are fibrillar muscles that power the flight through stretch activated asynchronous contractions allowing very high frequency oscillations, whereas the DFM are tubular muscle fibers with a cross-striated pattern that adjust the angle and rotation of the wings through contraction stimulated by nerve impulse.

Although later stages of muscle formation have been studied in great details, the understanding of early events in myoblast differentiation in the adepithelial layer of the wing disc is limited. One challenge is that the pool of myoblasts is inherently heterogeneous, as cells undergo differentiation and dynamically communicate with the epithelial layer through ligand receptor interactions (*7*, *8*). The advancement of single cell RNA-sequencing helped to begin resolving the heterogeneity of the AMPs (*7*, *9*, *10*). These cells share some of the features of the vertebrate adult muscle stem cells (*11*). The scRNA-seq studies revealed that myoblasts consist of several distinct populations of undifferentiated and differentiated cells and uncovered transcriptional programs underlying diversification of the IFM and DFM at these early stages (*9*). Furthermore, single cell experiments have identified genes with unique expression profiles that defined cell clustering, but were not previously associated with muscle development. Whether these genes are merely markers of these cell clusters or whether they may play a role in muscle development remains an open question for most genes. Interestingly, muscle specific knockdown of one of these genes, *Arginine kinase 1* (*Argk1*), resulted in pupal lethality suggesting that, at least some of them, may have an important function during muscle development.

*Argk1* belongs to the family of phosphagen kinases and is an analogue of vertebrate creatine kinase. It catalyzes the reversible transfer of the phosphate from adenosine triphosphate (ATP) to Arginine, and creates phosphoarginine, a high energy molecule that can be used to rapidly generate ATP, which is particularly relevant in tissues with high or fluctuating energy demand, such as muscle, neurons and spermatozoa (*12–14*). Indeed, dysfunctional creatine kinase has been associated with muscular dystrophies, inflammatory myopathies and metabolic disorders. In *Drosophila*, knockdown of Argk1 in motor neurons led to a rapid decline of ATP during axon burst firing (*15*). Even though extensive research has examined how phosphagen kinases maintaining energy homeostasis during different types of stress (*16*), their functions beyond these conditions are not well understood, and little is known about their role during development.

Here, we show that Argk1 is important in *Drosophila* flight muscles development. In proliferating myoblasts, Argk1 is required for cell growth, while at later stages it is needed for muscle growth and myofibril assembly. Argk1 knockdown results in significant disruption in energy homeostasis, manifested by an inability to maintain ATP levels and NAD+/NADH ratio, as well as aberrant mitochondrial morphology and changes in the expression of metabolic-related genes. Interestingly, developing Argk1-depleted muscles fail to undergo spontaneous muscle contractions, that is necessary for myofibril assembly during pupal stage and therefore may explain defects in the maturation of sarcomere and mitochondria. Thus, our work reveals a novel function of Argk1 during muscle development through its regulation of energy metabolism.

## Results

### Argk1 is expressed throughout the development of both direct and indirect flight muscles

We began by establishing the expression pattern of the Argk1 throughout muscle development using a GFP trap line *Argk1*^CB03789^ (Argk1::GFP). In this line, the GFP exon is fused in the appropriate reading frame to the endogenous Argk1 protein, thus, tagging Argk1 with GFP. This line was previously used to examine the expression of Argk1 *in vivo* (*9*, *15*). The wing imaginal discs harboring the myoblasts in the adepithelial layer were dissected from the *Argk*^CB03789^ third instar larva and stained with the myoblasts markers, Cut (Ct) and Twist (Twi). Confocal microscopy reveals strong GFP expression in most of Ct and Twi positive myoblasts and a weaker GFP signal in the epithelial cells of the wing disc (Figure 1A). This is in agreement with published scRNA-seq datasets showing Argk1 expression in myoblasts giving rise to IFM and DFM (*7*, *9*). To confirm that GFP accurately reflects the endogenous expression of Argk1, we performed RNAi experiments to deplete Argk1 in myoblasts using the *Mef2-GAL4* driver and two independent *UAS-Argk1-RNAi* transgenes: *JF02699* and *GD10436*. In both cases, the GFP signal was lost specifically in myoblasts labeled with anti-Ct antibody (yellow arrowhead), while the GFP expression was unaffected in the epithelial cells of the wing disc (Ct-negative cells, yellow asterisk) (Figure 1B). This is consistent with the experiments in adult brain where *GD10436* was shown to efficiently deplete Argk1 (*17*). Thus, we concluded that Argk1::GFP accurately reflects the endogenous expression of Argk1.

**Figure 1.**
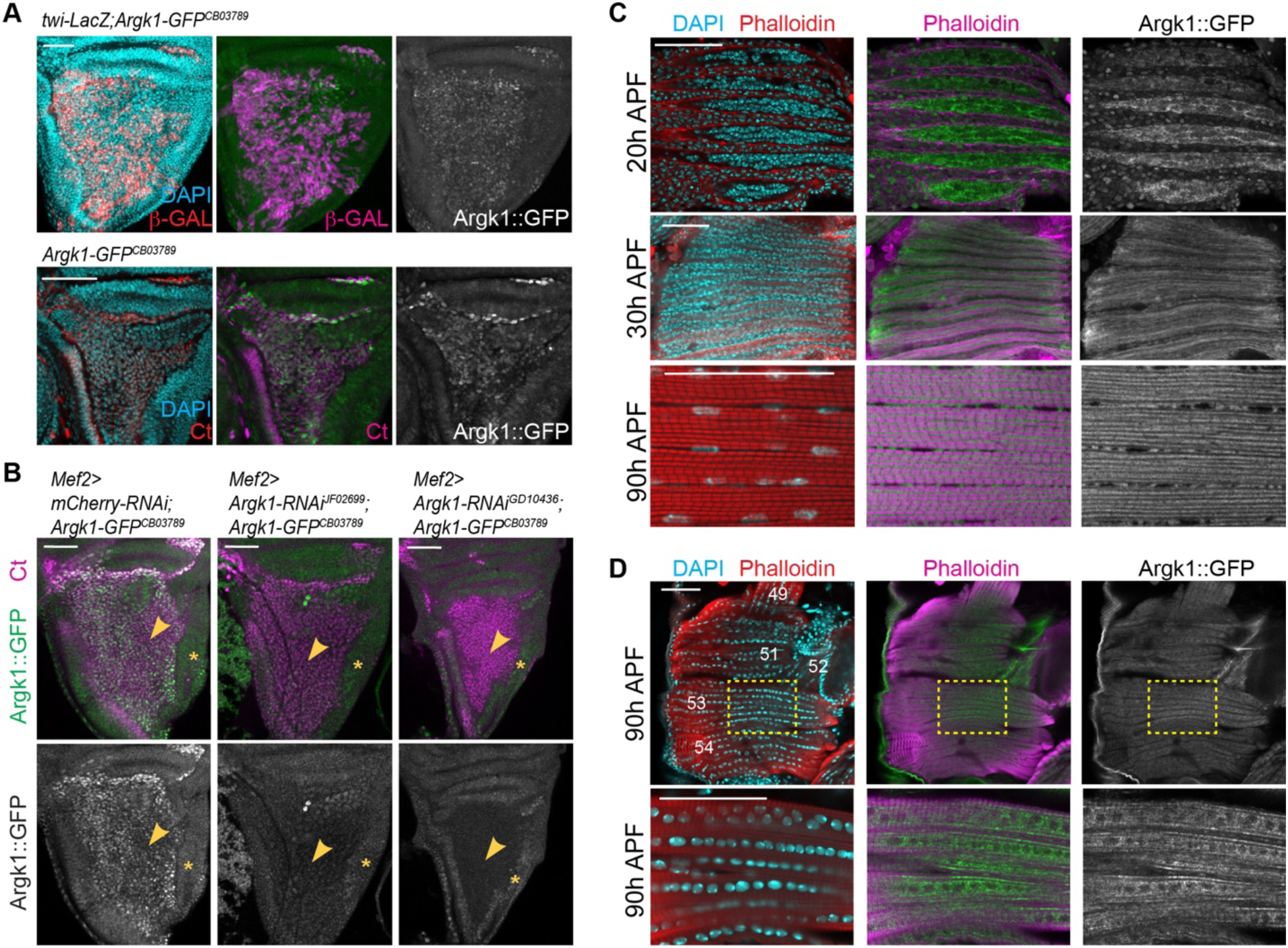
Argk1 is expressed throughout flight muscle development. (**A-D**) The GFP trap line *Argk1*^CB03789^ was used to track the expression of Argk1 (Argk1::GFP). (**A**, top panel) Maximum projection of z-stack from confocal images of wing discs of third instar larva stained with *Twi-LacZ* (β-gal) to label the myoblasts (magenta), confocal single plane image of (**B**, bottom panel) third instar larval wing discs stained with anti-Ct to label the myoblasts (magenta), (**C**) forming DLM-IFM at 20 h, 30 h, and sagittal sections of IFM at 90 h APF, and (**D**) DFM at 90 h APF stained with Phalloidin to mark the myofibers (red and magenta) and with DAPI to label the nuclei (cyan). The different muscles of DFM were noted, the yellow- dashed box indicates magnified area for the DFM_53 in bottom panel. Anterior to the left. (**B**) Argk::GFP expression is lost in the myoblasts as indicated by the yellow arrowhead in both *Mef2>Argk1-RNAi* lines, while expression persisted in epithelial cells (yellow asterisk). Genotypes are: (**A**) w; *twi-LacZ*/+; *P{PTT-GB}Argk1^CB03789^*/+ *w*; +; *P{PTT-GB}Argk1^CB03789^*/+ (**B**) w; +; *Mef2-GAL4*, *P{PTT-GB}Argk1^CB03789^* /*UAS-mCherry-RNAi* *w; +; Mef2-GAL4*, *P{PTT-GB}Argk1^CB03789^*/*UAS-Argk1-RNAi^JF02699^* *w; +; Mef2-GAL4*, *P{PTT-GB}Argk1^CB03789^*/*UAS-Argk1-RNAi ^GD10436^* (**C-D**) *w*; +; *P{PTT-GB}Argk1^CB03789^*/+ Scale 50 μm.

Next, we assessed Argk1 expression throughout flight muscle development. During pupal stages, myoblasts migrate and fuse to form multinucleated myotubes and then undergo myofibrillogenesis and growth. DLMs were dissected at 20 h, 30 h and 96 h after pupa formation (APF) representing distinct timepoints in myogenesis. Tissues were stained with Phalloidin and DAPI to visualize developing muscles and nuclei, respectively. We found that Argk1::GFP is strongly expressed in both fusing myoblasts and multinucleated myotubes at 20 h APF, near the immature myofibrils in the myofibers at 30 h APF, and in between myofibrils in the mature muscles at 96 h APF (Figure 1C). Since Argk1 is expressed in the myoblasts that give rise to both IFM and DFM (Figure 1A) (*9*), we assessed the expression of Argk1 in mature DFM as well. The muscles were dissected from pharate-staged Argk1::GFP animals at 90 h APF and the GFP signal was examined. Sagittal sections of DFM 51, DFM 53 and DFM 54 showed strong GFP fluorescence in these muscles while nuclei were largely GFP negative (Figure 1D). Thus, Argk1 is highly expressed in proliferating myoblasts, and its expression persists in myofibers of both IFM and DFM throughout muscle development.

### Argk1 is required in adult muscles for animal viability

We used a panel of well-established muscle-specific GAL4 drivers that are expressed at different time points and in different muscles to explore the requirement for Argk1 in muscle development. The depletion of Argk1 with the pan-muscle *Mef2-GAL4* driver, which is expressed throughout development, using the *UAS-Argk1-RNAi* line *JF02699* led to lethality at pharate stage (Figure 2A). The knockdown of Argk1 using an independent *UAS-Argk1-RNAi* transgene *GD10436* also resulted in lethality, albeit at a later time point, as most animals died within 3 days post-eclosion (Figure 2A), which is in agreement with previous reports (*18*). Unlike *Mef2-GAL4*, the *Act88F-GAL4* driver is activated specifically in the IFM at the onset of myotube differentiation around 20 h APF (*19*). The knockdown of Argk1 using *JF02699* under the control of *Act88F-GAL4* resulted in predominantly male lethality within five days post-eclosion (Figure 2B). Although *GD10436* did not result in lethality with *Act88F-GAL4*, the muscles of these animals were dysfunctional as revealed by the flight test. While the control animals landed at the top of the cylinder, almost half of *GD10436* and three quarters of *JF02699* flies failed to reach the top and landed at the bottom (Figure 2C), thus indicating that flight muscles are not fully functional.

**Figure 2.**
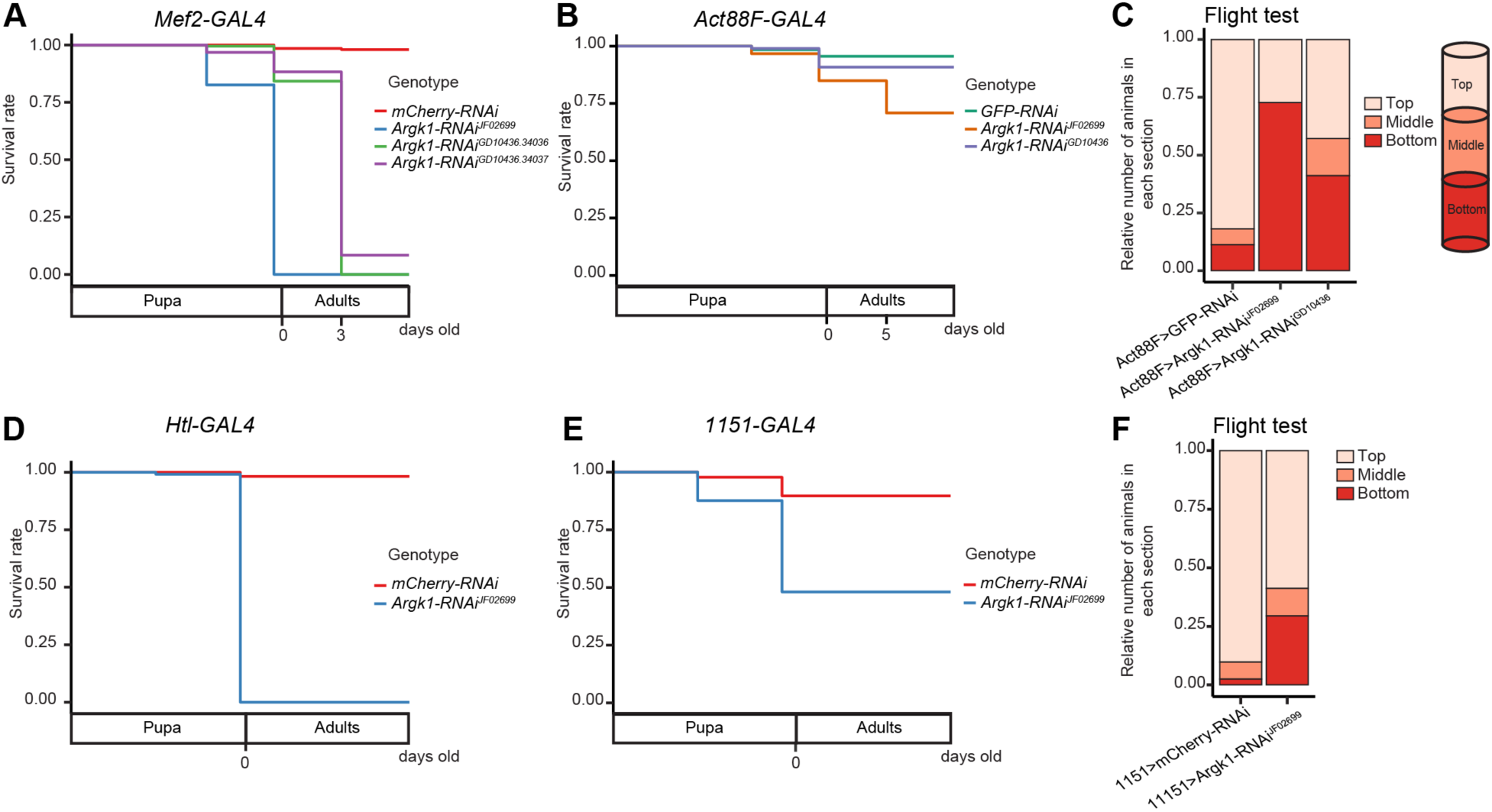
The loss of Argk1 specifically in muscle results in increased lethality during development and flightless animals. (**A-B**, **D-E**) Kaplan-Meier-like plot to illustrate survival rate of animals through late stages of development, only pupa and adult animals were quantified. The expression of Argk1 was knock down using three *UAS-RNAi* lines: one transgene *JF02699* and two *GD10436* (stocks 34036 and 34037 with different insertion sites) and compared to control *UAS-mCherry-RNAi* or *UAS-GFP-RNAi*. The drivers used are (**A**) *Mef2-GAL4*, (**B**) *Act88F-GAL4*, (**C**) *Htl-GAL4* and (**D**) *1151-GAL4*. Data represent mean of survival rate at each developmental stage scored. (**A**) n= 60 flies per genotype, (**B**) n= 90 flies per genotype, (**D**) n= 120 flies per genotype, and (**E**) n= 300 flies per genotype. Experiment was repeated N= 2-8 times. (**C-F**) Flight ability scored by quantifying the percentage of flies landing on section of the cylinder (top, middle and bottom). Stacked bars, (**C**) n = 11-116 flies /genotype, (**F**) n = 17-41 flies /genotype. The ability to flight of the animals that survived in **B** and **E** were assessed in **C** and **F**, respectively. Most males *Act88F>Argk1-RNAi^JF02699^* and *1151>Argk1-RNAi^JF02699^* died, and only 11 and 17, respectively, survived for the flight ability test. Genotypes are: (**A**) w; +; *Mef2-GAL4*/*UAS-mCherry-RNAi* w; +; *Mef2-GAL4*/*UAS-Argk1-RNAi^JF02699^* w; +; *Mef2-GAL4*/*UAS-Argk1-RNAi ^GD10436.34036^* w; +; *Mef2-GAL4*/*UAS-Argk1-RNAi ^GD10436.34037^* (**B-C**) *Act88F-GAL4*; *UAS-GFP-RNAi/+*; + *Act88F-GAL4*; +; *UAS-Argk1-RNAi^JF02699^/+* *Act88F-GAL4*; +; *UAS-Argk1-RNAi ^GD10436.34036^/+* (**D**) w; +; *Htl-GAL4*/*UAS-mCherry-RNAi* w; +; *Htl-GAL4*/*UAS-Argk1-RNAi^JF02699^* (**E-F**) *1151-GAL4*; +; *UAS-mCherry-RNAi/+* *1151-GAL4*; +; *UAS-Argk1-RNAi^JF02699^/+*

The *UAS-Argk1-RNAi* transgene *JF02699* that gave stronger phenotypes than *GD10436* was further tested with two additional GAL4 drivers, *Htl-GAL4* and *1151-GAL4*. The *Htl-GAL4* is expressed in the mesoderm and muscle lineages including adult muscle precursors (*7*, *20*). Depletion of Argk1 with *Htl-GAL4* resulted in lethality at the pharate stage (Figure 2D), similar to what we found with *Mef2-GAL4*. The other GAL4 driver, *1151-GAL4*, is expressed in myoblasts giving rise to adult skeletal muscles, but it is not expressed in larval muscle lineage (*5*). The knockdown of Argk1 with *1151-GAL4* resulted in partial lethality as half of the animals died at pharate stage (Figure 2E). Notably, the surviving animals showed defective muscle function, since more than the third of flies failed to reach the top of the cylinder in the flight test (Figure 2F).

We concluded that Argk1 is essential throughout muscle development, as its knockdown using pan-muscle or late muscle specific GAL4 drivers resulted in either lethality or decreased muscle function, respectively.

### Argk is required for the development of both direct and indirect flight muscles

The results of knocking down Argk1 expression with a panel of GAL4 drivers suggest that Argk1 plays an important role in the muscle. To determine how the loss of Argk1 function affects flight muscle development, we examined the structure of DFM and IFM in animals expressing the *UAS-Argk1-RNAi ^JF02699^* transgene under the control of *Mef2-GAL4* (*Mef2>Argk1-RNAi*). Transverse and sagittal sections of *Mef2>Argk1-RNAi* and control pharate adults were stained with Phalloidin and anti β-PS-integrin antibody to visualize muscle structure (Figure 3A-B). Notably, the *Argk1*-depleted DFM and IFM were significantly reduced in size. Quantifications confirmed that the cross-sectional area of DLM 3 and DLM4 were statistically significantly smaller than in wild type control (Figure 3C). These results were validated with another *UAS-Argk1-RNAi* line *GD10436* that produced similar, but milder phenotype (Figure S1A-B). The milder phenotype induced by *GD10436* was consistent with the results shown above (Figure 2A), thus indicating that the line *JF02699* was stronger than *GD10436*.

**Figure 3.**
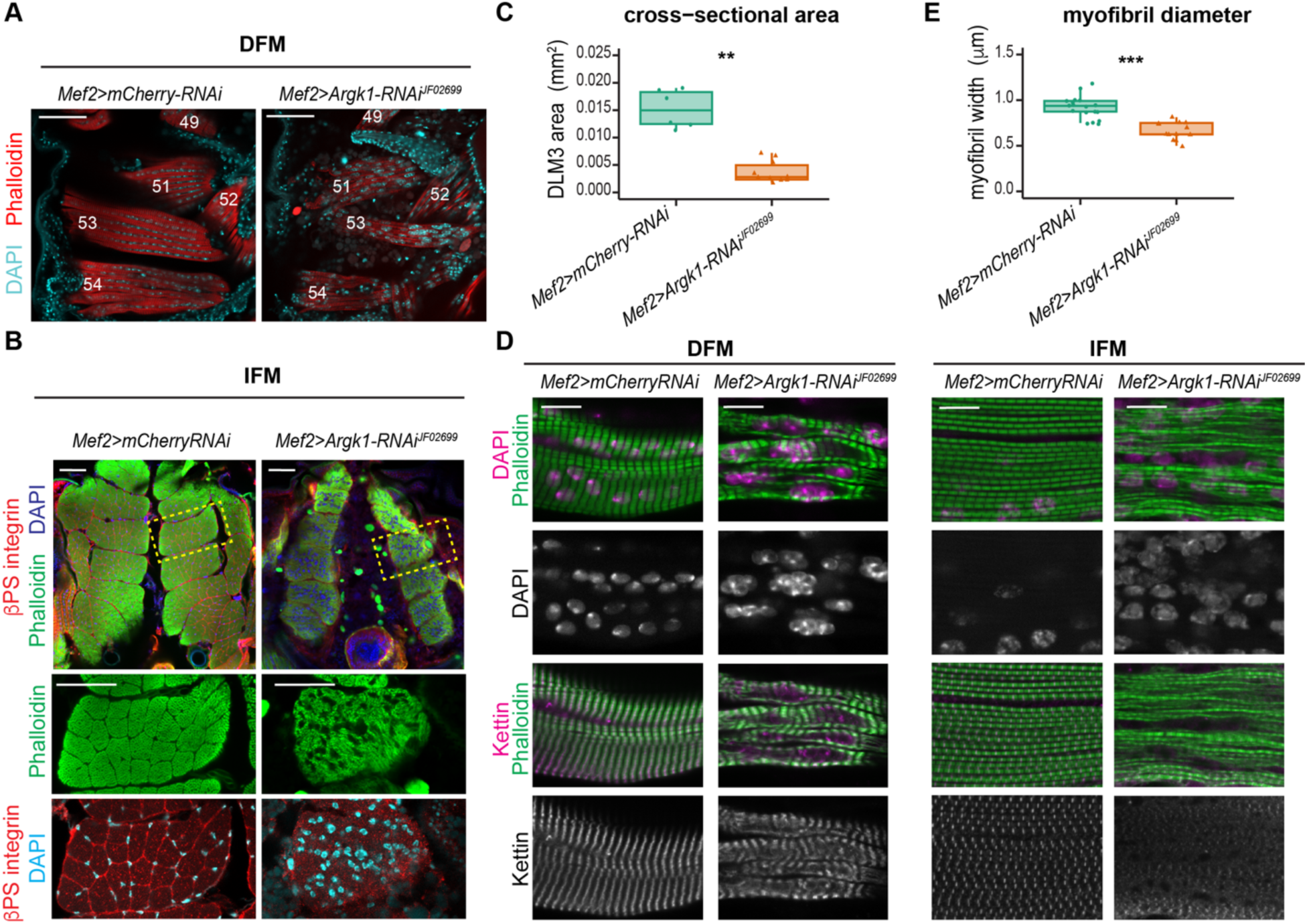
Argk1 knockdown with *Mef2-GAL4* results in severe defects in both IFM and DFM. (**A-D**) The structure of the DFM and IFM examined in *Mef2>Argk1-RNAi^JF02699^* and control *Mef2>mCherry-RNAi* animals staged at 90h APF. (**A**) Confocal single plane images of DFM in a sagittal view. Hemithorax sections stained with Phalloidin (red) and DAPI. DFM are numbered in white as in (*38*). Anterior is to the left. (**B**) Confocal single plane images of IFM in a transverse view of the dorsal longitudinal muscles (DLM). Sections stained with Phalloidin (green), DAPI (blue) and βPS-integrin. Yellow-dashed box indicates magnified area for DLM3 (bottom panel). Dorsal is up. (**C**) Quantification of the cross-sectional area for the DLM3 per animal. N = 6-10 thoraces/genotype, Wilcoxon rank-sum test, ** p<0.01. (**D**) Confocal single plane images of DFM-53 (left panel) and IFM-DLM (right panel) in a sagittal view. Hemithorax sections stained with Phalloidin, DAPI and anti-Kettin. (**E**) Quantification of myofibril diameter for the IFM-DLM per animal. N = 13-18 thoraces/genotype, Welch Two Sample t-test, *** p<0.001. All box plot shows median (middle), interquartile range (box), 1.5x the interquartile range (whiskers), and individual data (points). Scale (**A-B**) 50 μm, and (D) 10 μm. Genotypes are: w; +; *Mef2-GAL4*/*UAS-mCherry-RNAi* w; +; *Mef2-GAL4*/*UAS-Argk1-RNAi^JF02699^*

To visualize the overall sarcomere organization of Argk1-depleted muscles, sagittal sections of DFM and IFM were stained with Phalloidin and anti-Kettin antibody to label the Z-discs. Severe defects in myofibril assembly were found in both DFM and IFM of *Mef2>Argk1-RNAi* pharate adults (Figure 3D), resulting in a significant reduction in myofibril diameter (Figure 3E), thus suggesting a defect in myofibrillogenesis (*21*). The phenotypes were confirmed with the other *UAS-Argk1-RNAi* line *GD10436* (Figure S1C) by quantifying defects in both sarcomere length and myofibril diameter (Figure S1D-E).

Overall, our findings further support the requirement of Argk1 during the development of both flight muscles, DFM and IFM, for muscle growth and myofibrillogenesis.

### Argk1 is required for flight muscle growth at late stages of development

To determine whether defects in muscle growth and myofibrillogenesis are due to the requirement of Argk1 in late muscle development, Argk1 was depleted using the late muscle driver *Act88F-GAL4*, whose expression spans from the onset of myotube differentiation (∼20 h APF) to muscle maturation (*19*). The transverse sections of thoraces from 5-days old adults stained with Phalloidin and anti-β-PS-integrin antibodies revealed a significant reduction in the IFM cross-sectional area, as quantified for DLM4 (dashed yellow box), in *Act88F>Argk1-RNAi ^JF02699^* compared to control flies (Figure 4 A-B). Moreover, sagittal sections stained with Phalloidin and anti-Kettin antibodies revealed reduced myofibril diameter in Argk1-depleted muscles albeit sarcomere length was normal (Figure 4 C-E). Thus, Argk1 is required in late muscle development for muscle growth and for myofibril and sarcomere assembly.

**Figure 4.**
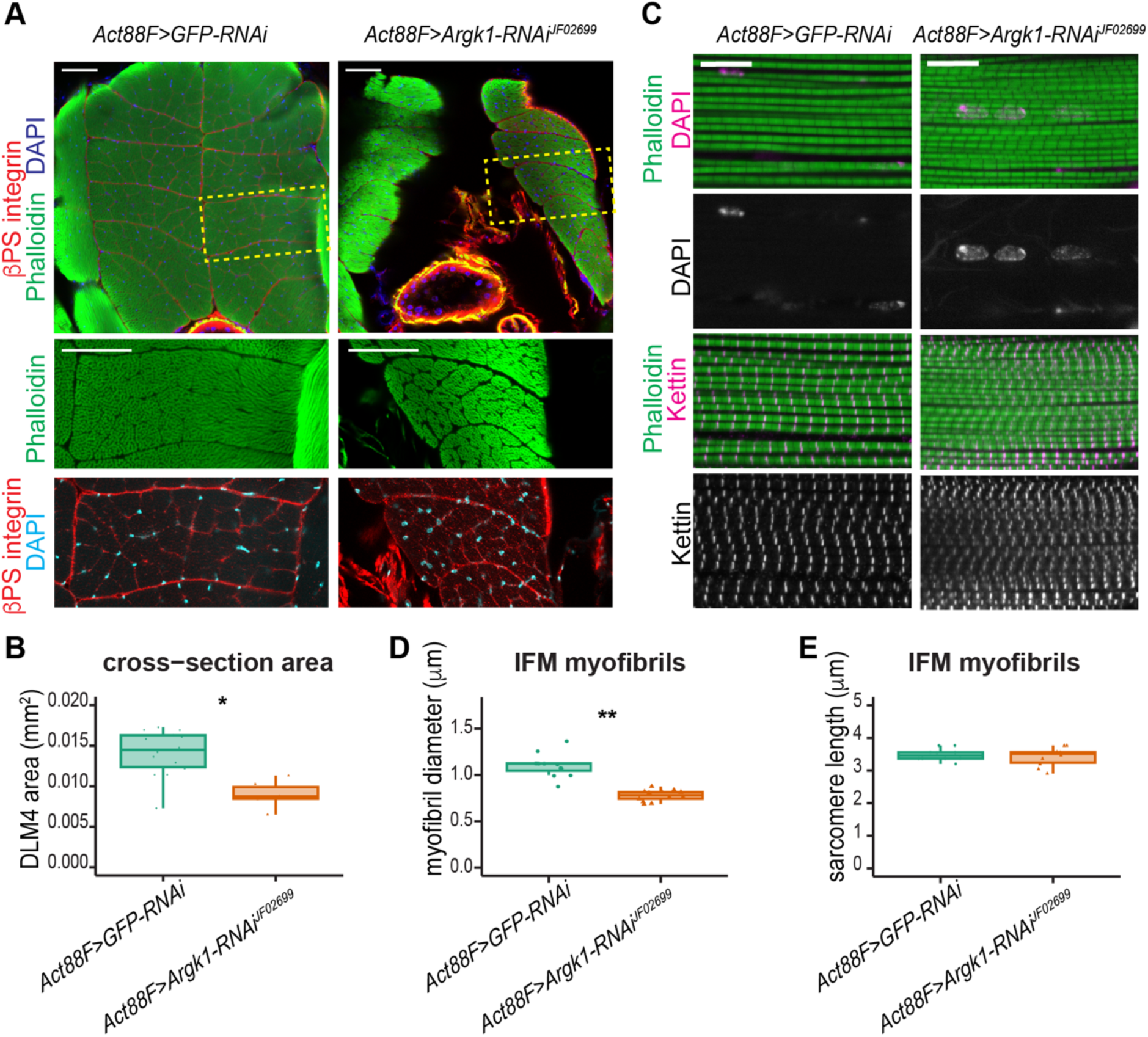
Argk1 knockdown with *Act88F-GAL4* results in severe muscle defects in IFM. (**A-E**) The structure of the IFM examined in 1-2 days old *Act88F>Argk1-RNAi^JF02699^* and control *Act88F>GFP-RNAi* animals. (**A**) Confocal single plane images of IFM in a transverse view of the dorsal longitudinal muscles (DLM). Sections stained with Phalloidin (green), DAPI (blue) and βPS-integrin. Yellow-dashed box indicates magnified area for DLM4 (bottom panel). Dorsal up. (**B**) Quantification of the cross-sectional area for the DLM4 per animal. N = 6-14 thoraces/genotype, Wilcoxon rank-sum test, * p<0.05. (**C**) Confocal single plane images of IFM-DLM in a sagittal view. Hemithorax sections stained with Phalloidin, DAPI and anti-Kettin. (**D**) Quantification of myofibril diameter for the IFM-DLM per animal. N = 12 thoraces/genotype, Wilcoxon rank-sum test, ** p<0.01. (**E**) Quantification of sarcomere length for the IFM-DLM per animal. N = 10 thoraces/genotype, Wilcoxon rank-sum test, p>0.05. All box plot shows median (middle), interquartile range (box), 1.5x the interquartile range (whiskers), and individual data (points). Scale (**A**) 50 μm, and (**C**) 10 μm. Genotypes are: *Act88F-GAL4; +; UAS-mCherry-RNAi/+* *Act88F-GAL4*; +; *UAS-Argk1-RNAi^JF02699^/+*

### Argk1 impacts cell size of myoblasts at the onset of flight muscle development

Next, we investigated whether Argk1 has a role at the early stages of muscle development, specifically during myoblast proliferation. Examining overall distribution of myoblasts in the *Mef2>Argk1-RNAi* wing discs by staining with both anti-Cut antibodies and *Htl-lexA>lexO-CD8-GFP* (Patel et al. 2022) did not reveal severe changes compared to the control (Figure 5A). However, we noticed a reduction in the depth of the layer of Argk1-depleted myoblasts normalized to the depth of the disc proper compared to control (Figure 5B-C). The reduced depth of the myoblast layer could be due to a reduction in either the number of cells and/or in the cell size. To examine the former possibility, we determined the number of myoblasts per area. The myoblasts nuclei were labeled with *Htl>nls-RFP*, and both the number of DAPI and RFP positive nuclei per image were counted in an automated manner using the ImageJ software (Figure 5D). No significant differences were observed between *Mef2>Argk1-RNAi* and the control (Figure 5E-F). In addition, we assessed cell proliferation by staining wing imaginal discs with a mitotic marker, anti-phospho-histone 3 (pH3) antibody. To distinguish the pH3 positive myoblasts from pH3 positive epithelial cells, *twi-LacZ* was used to label the myoblasts (Figure 5G). The number of pH3-positive myoblasts relative to the number of cells per image did not show significant change between Argk1-depleted and control myoblasts (Figure 5H).

**Figure 5.**
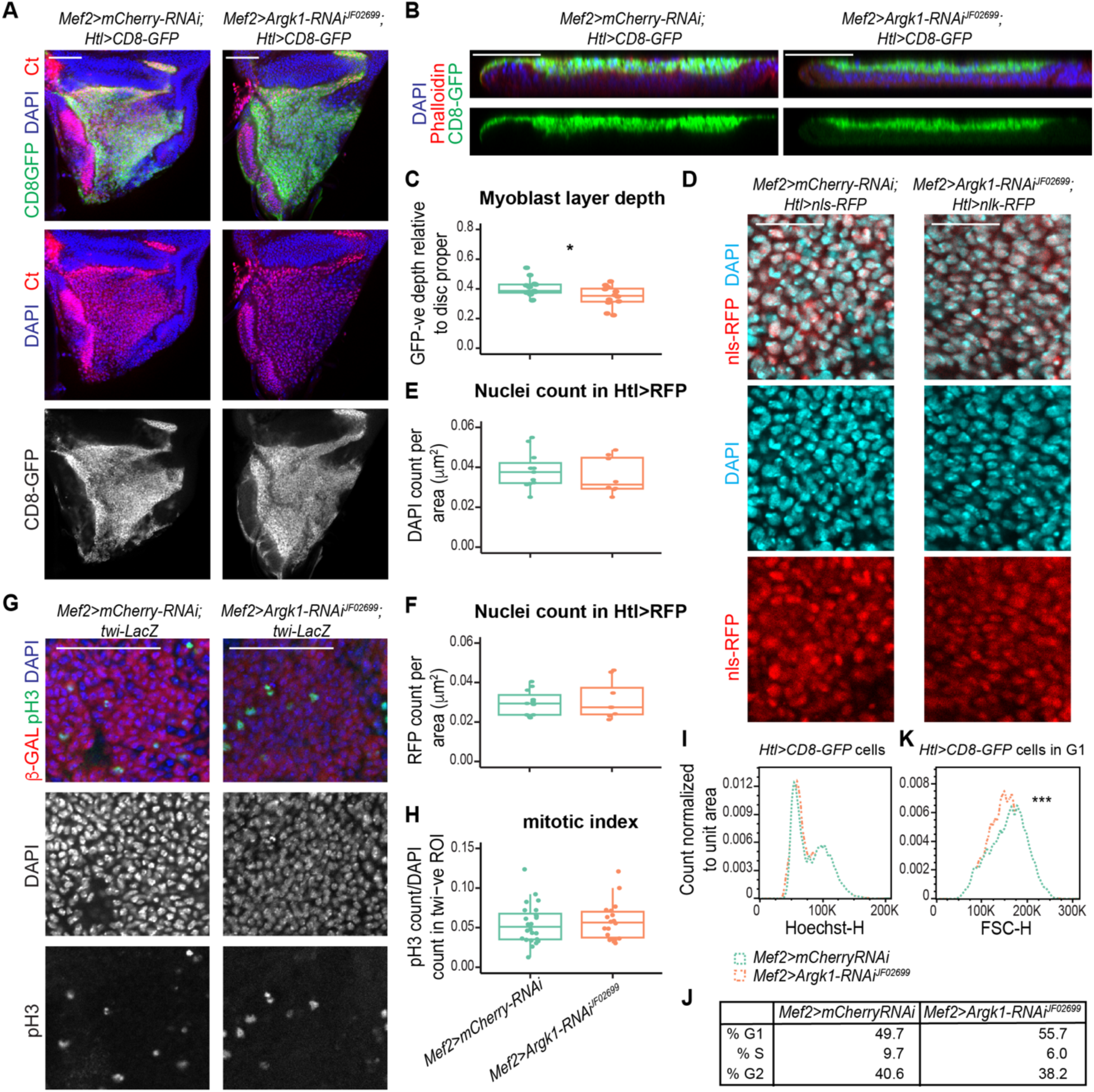
Knockdown of Argk1 reduces cell size but does not affect cell cycle progression. The myoblasts in the adepithelial layer of the third instar larva wing discs compared between *Mef2>Argk1-RNAi^JF02699^* and the control *Mef2>mCherry-RNAi*. (**A**) Confocal single plane images of wing discs stained with anti-Ct (red), *Htl-LexA*; *lexO-CD8-GFP* (green) to label the myoblasts and DAPI (blue). (**B**) XZ orthogonal view wing discs centered in the notum stained with phalloidin (red), *Htl-LexA*; *lexO-CD8-GFP* (green) and DAPI (blue). (**C**) Quantification of the depth of the myoblast layer from XZ orthogonal view in B. Ratio between depth of the GFP positive cells relative to whole disc per disc. n = 12-13 discs per genotype, two-tailed Student’s t- test, p<0.05. N=2 independent experiments. (**D**) Confocal single plane images of wing discs labeled with *Htl-LexA*; *LexO-nls-RFP* (red) to mark the myoblasts and DAPI (blue). (**E-F**) Quantification of the number of myoblasts per area using (**E**) DAPI and (**F**) *Htl>nls-RFP*. n = 8-11 discs/genotype, Wilcoxon rank-sum test, p<0.05. N=3 independent experiments. (**G**) Confocal single plane images of wing discs stained with anti-pH3 (green) to label mitotic cells, *twi-LacZ* (red) to mark the myoblasts and DAPI (blue). (**H**) Quantification of the number of pH3 positive cells relative to the number of nuclei (DAPI count) within the Twist positive area. n = 18-24 discs/genotype, Wilcoxon rank-sum test, p<0.05. N=3 independent experiments. (**I**) Histogram of DNA content vs counts for myoblasts sorted using *Htl>CD8-GFP* cells and normalized to unit area by flow cytometry, n=7168 and 7894 cells/genotype. N=2 independent experiments. (**J**) Percent of cells in G1, S and G2 from (**I**) as determined by Dean-Jett-Fox model. (**K**) Histogram of relative cell size as measured by forward scatter vs counts for myoblast cells selected in G1 from (**I**) normalized to unit area, n=3403 and 4282 cells per genotype. Welch Two Sample t-test, *** p<0.001. N=2 independent experiments. All box plot shows median (middle), interquartile range (box), 1.5x the interquartile range (whiskers), and individual data (points). Scale (**A,B,G**) 50 μm, and (**D**) 20 μm. Genotypes are: (**A-C**, **I-K**) w; *lexO-CD8-GFP/+*; *Htl-LexA, Mef2-GAL4*/*UAS-mCherry-RNAi* w; *lexO-CD8-GFP/+*; *Htl-LexA, Mef2-GAL4*/*UAS-Argk1-RNAi^JF02699^* (**D-F**) w; *lexO-nls-RFP/+*; *Htl-LexA, Mef2-GAL4*/*UAS-mCherry-RNAi* w; *lexO-nls-RFP/+*; *Htl-LexA*, *Mef2-GAL4*/*UAS-Argk1-RNAi^JF02699^* (**G-H**) w; *twi-LacZ/+*; *Mef2-GAL4*/*UAS-mCherry-RNAi* w; *twi-LacZ/+*; *Mef2-GAL4*/*UAS-Argk1-RNAi^JF02699^*

To quantify cell cycle progression along with cell size of the Argk1-depleted myoblasts, we performed flow cytometry experiments. The wing discs from *Mef2>mCherry-RNAi Htl>CD8-GFP* and *Mef2>Argk1-RNAi Htl>CD8-GFP* animals were dissociated into single cell suspension, stained with Hoechst and sorted by Fluorescence-Activated Cell Sorting (FACS) to determine DNA content and forward light scatter (FSC), a measurement of cell size. The myoblasts were distinguished from epithelial cells of the wing disc by the presence of GFP while their cell cycle profile was determined based on the intensity of Hoechst fluorescence that reflects the DNA content (Figure 5I). The Dean-Jett-Fox model was fitted to estimate the percentage of G1 (or 2C), S and G2 (4C) cells for the myoblasts. The percentage of G1, S and G2 cells were comparable between control and Argk1-depleted myoblasts (Figure 5J), which is consistent with normal mitotic index in *Mef2>Argk1-RNAi,* as determined by immunofluorescence (Figure 5G-H). However, the cell size of *Mef2>Argk1-RNAi* myoblasts was significantly smaller than that of control cells as revealed by the measurement of forward scatter (FSC) of the G1 cells (Figure 5K). These results suggest that although Argk1-depleted myoblasts proliferate normally, their size is smaller, which may explain the reduction in the depth of the myoblasts layer in the *Mef2>Argk1-RNAi* wing imaginal discs. These findings support the idea that Argk1 is important for cell growth at the early onset of flight muscle development, but it is not required for cell cycle progression.

### Single cell RNA-sequencing identifies new myoblast markers

Transcriptional changes orchestrate and drive the morphological transitions that occur during muscle development (*21*). Given that knocking down Argk1 impacts muscle development, we aimed to determine how the transcriptome changes in Argk1-depleted myoblasts. However, the physical association of the myoblasts with the epithelial cells of the wing disc makes bulk approaches unsuitable for their profiling. Therefore, we performed single cell RNA-sequencing (scRNA-seq) to resolve the myoblast-specific changes upon Argk1 depletion without the confounding contribution from the epithelial cells. The wing discs of *Mef2>Argk1-RNAi* and *Mef2>mCherry-RNAi* third instar larva were dissociated into single cell suspension and the transcriptomes of individual cells were determined using the 3’ Gene Expression assay (10X Genomics). The data were processed using Seurat v5 package (*22*) to cluster cells based on gene expression similarity. After filtering poor quality cells and applying Principal Component Analysis, the non-linear dimensionality reduction algorithm, UMAP, was used for visualization of cell clusters. Epithelial cells, tracheal cells and myoblasts were identified based on the expression of the respective markers: *Fasciclin 3* (*Fas3*) and *grainy head* (*grh*) for epithelia, *trachealess* (*trh*) for trachea, and *Zn finger domain 1* (*zfh1*), *Holes in muscles* (*Him*) and *Secreted protein, acidic, cysteine-rich* (*SPARC*) for myoblast. A total of 11,489 myoblasts across two genotypes and two replicates each were selected for downstream analysis (Figure 6A, Data S1). Based on the differential expression of canonical DFM and IFM marker genes, *Ct* and *vg*, we identified three clusters that will give rise to DFM, and eight clusters that will give rise to IFM (Figure 6A-B).

**Figure 6.**
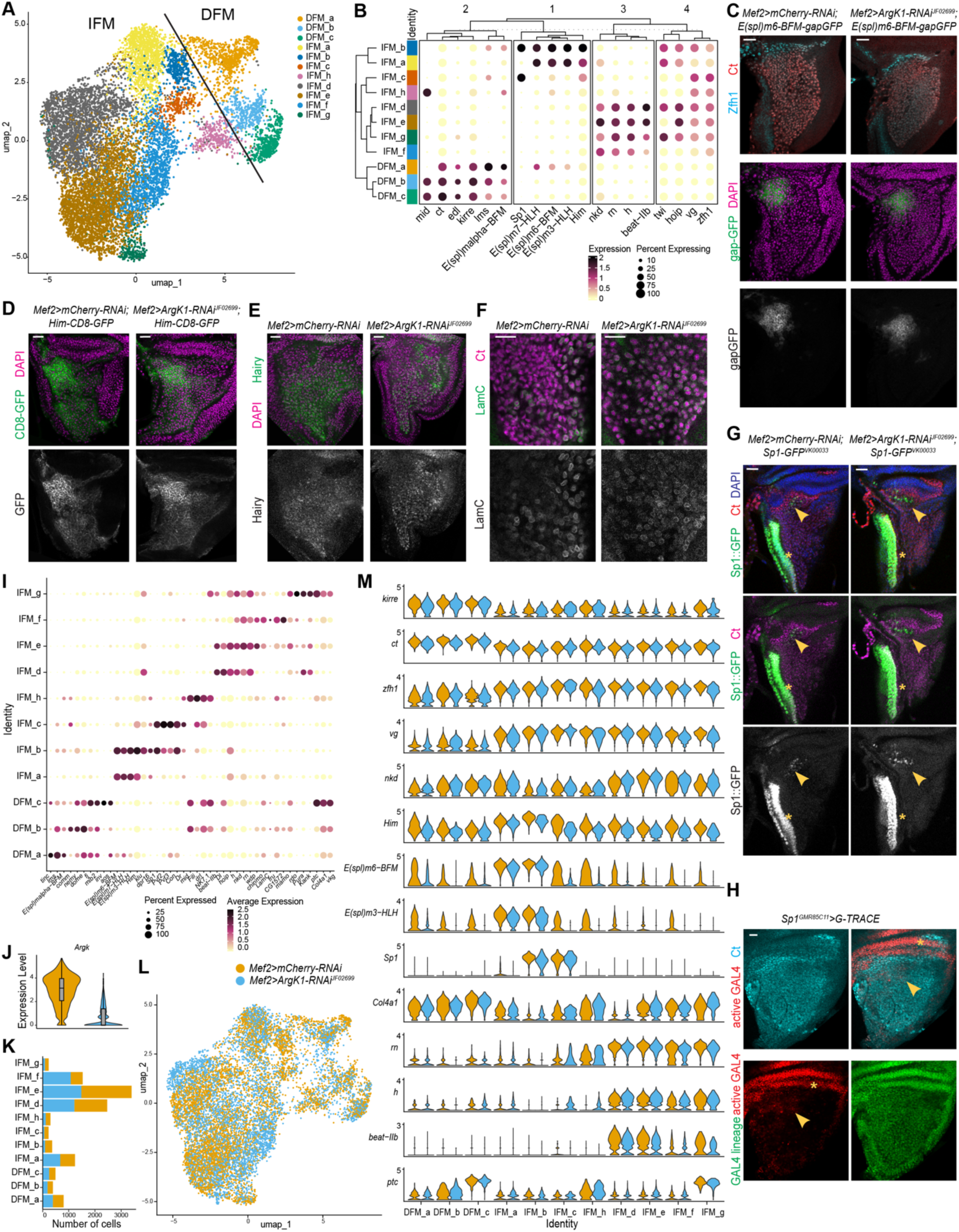
Knockdown of Argk1 does not alter the overall identity of the undifferentiated and differentiated myoblasts. (**A**) Two-dimensional UMAP representation of both the combined *Mef2>Argk1-RNAi^JF02699^* (5,681 myoblasts) and *Mef2>mCherry-RNAi* (5,808 myoblasts) from third instar larval wing discs datasets, colored by (A) cell cluster and (L) genotype. (**B**) Hierarchical dot plots showing the expression levels of the top known markers for myoblasts. Hierarchical clustering by k-means to group both the clusters of cells and the genes into groups based on similarity. Color intensity shows the average expression level normalized and scaled. The size of the dot represents the fraction of cells within a cluster expressing each gene (**C-G**) Confocal single plane images of *Mef2>mCherry-RNAi* and *Mef2>Argk-* RNAi^JF02699^ third instar larval wing discs stained with (**C**) anti-Ct (red), anti-Zfh1 (cyan), and labeled with *E(pl)m6-BFM-gapGFP* (GFP) and DAPI (magenta), (**D**) *Him-CD8-GFP* (green) and DAPI (magenta), (**E**) anti-Hairy (green) and DAPI (magenta), (**F**) anti-Ct (magenta) and anti-LamC (green), and anti-Ct (red/magenta), *Sp1-GFP* (green) and DAPI (blue). (**H**) Confocal single plane images of *Sp1^GMR85C11^>G-TRACE* third instar larval wing discs stained with anti-Ct (cyan) to label myoblasts, showing the lineage of *Sp1-GAL4* (green) and the active GAL4 (red). (**I**) Dot plots showing the expression levels of selected genes from the top markers list identified in this dataset. Color intensity shows the normalized and scaled average expression level. The size of the dot represents the fraction of cells within a cluster expressing each gene (**J**) Violin plot showing the distribution of expression of Argk1 for the combined datasets *Mef2>Argk1-RNAi^JF02699^* (blue) and *Mef2>mCherry-RNAi* (yellow). (**K**) Total number of cells per cluster per genotype relative to the total number of myoblasts per genotype of the combined datasets *Mef2>Argk1-RNAi^JF02699^* (blue) and *Mef2>mCherry-RNAi* (yellow). (**M**) Stacked violin plots representing the distribution of expression of known myoblast markers across clusters of the of the combined datasets. Genes include *kin of irre* (*kirre*), *cut* (*ct*), *Zn finger homeodomain 1* (*zfh1*), *vestigial* (*vg*), *naked cuticle* (*nkd*), *Holes in muscle* (*Him*), *Enhancer of split m6* (*E(spl)m6-BFM*), *Enchancer of split m3* (*E(spl)m3-HLH*), *Sp1*, *Collagen type IV alpha 1* (*Col4a1*), *rotund* (*rn*), *hairy* (*h*), *beaten path IIb* (*beat-IIb*), *patched* (*ptc*). Data were split by genotype, *Mef2>mCherry-RNAi* (yellow) a*nd Mef2>Argk-RNAi^JF02699^* (blue). Scale 20 μm. Genotypes are: w; +; *Mef2-GAL4*/*UAS-mCherry-RNAi* w; +; *Mef2-GAL4/UAS-Argk1-RNAi^JF02699^*

Before comparing the transcriptomes of wild type and Argk1-depleted myoblasts, we first annotated the cell clusters. We established the inherent hierarchical order of the cell clusters by k-means clustering using a panel of myoblast markers derived from ours and others scRNA-seq experiments (*7*, *9*) (Figure 6B, Data S2). Notably, all eight myoblast clusters giving rise to IFM were grouped together and were distinct from three DFM clusters (Figure 6B). As a result of hierarchical clustering, the markers fell into four groups: Group 1, Group 2, Group 3 and Group 4. The markers specific to IFM included *vg* and *zfh1* in Group 4, while the markers for DFM included *lms*, *kirre* and *ct* in Group 2. Genes expressed in the undifferentiated myoblasts, IFM_a, IFM_b, IFM_c and DFM_a, included the *E(spl)* genes, *Him* and *Sp1* in Group 1, and markers expressed in the differentiating myoblasts included *hairy* in Group 3. Thus, the analysis of hierarchical clustering defined the clusters of cells giving rise to distinct muscle types along with undifferentiated and differentiated myoblasts.

Next, we visualized the expression of some of these markers by immunofluorescence using antibodies and reporter lines (Figure 6C-G). The genes listed in Group 1, *E(spl)m6-BFM*, *Him* and *Sp1*, which are all markers of the undifferentiated IFM clusters, showed high expression levels in the adepithelial layer near the anterior region of the notum and the dorsal hinge (Figure 6C, D, G). Although *Him* is expressed throughout the myoblast layer, it showed significantly higher expression in the specified region (Figure 6B,D). The validated *Sp1-GFP^VK00033^* reporter (*23*) is expressed in a very restricted group of myoblasts (yellow arrowhead, Figure 6G) in addition to a stripe of epithelial cells (yellow asterisk). The gene *hairy* listed in Group 3 as a marker for the differentiating IFM clusters, was spatially restricted across the adepithelial layer (Figure 6B,E). These results are consistent with the observation that the positions of the undifferentiated and differentiated cell clusters are spatially restricted across the myoblasts in the adepithelial layer, as cell clusters containing undifferentiated cells are localized near the anterior region of the notum (*9*).

Since Sp1 has not been previously validated as a marker for clusters of undifferentiated myoblasts, we used gTRACE (*24*) to genetically trace the Sp1-expressing cells. In this method, GFP labels the cells that expressed Sp1 in the past (GAL4 lineage), while RFP marks the cells that expressed Sp1 at the time of analysis (active GAL4). We tested four lines *Sp1-GAL4^GMR85C11^*, *Sp1-GAL4^GMR85C01^*, *Sp1-GAL4^GMR85C05^* and *Sp1-GAL4^GMR85C10^* that reflect the activity of the corresponding enhancers upstream *Sp1* gene. Among them, only *Sp1-GAL4^GMR85C11^* was expressed in myoblasts (yellow arrowheads, Figure 6H), in addition, to the expression in the epithelial cells (yellow asterisks, Figure 6H). Notably, the myoblast specific expression of *Sp1-GAL4^GMR85C11^* largely matched the pattern of expression of the *Sp1-GFP^VK00033^* reporter (yellow arrowheads, Figure 6G-H). In contrast, GFP, labeling the GAL4 lineage, marked all myoblasts as visualized by counterstaining with anti-Ct antibody (Figure 6H), thus suggesting that Sp1-expressing cells in IFM_b and IFM_c clusters are likely the progenitor cells for differentiating myoblasts. This conclusion is in concordance with tracing experiments for two other genes from Group 1: *E(spl)m3-HLH* and *E(spl)m6-BFM* that also mark progenitor cells (*9*).

To further explore the transcriptional differences between the cell clusters, the expression of top 30 cluster specific marker genes was visualized by a hierarchical clustered heatmap (Figure S2A, Data S3, Data S4). In total, eight gene signatures were identified by k-means clustering across the cell clusters. Concordantly, the IFM clusters containing undifferentiated myoblasts (IFM_a, IFM_b, IFM_c) were clustered separately from the IFM clusters containing differentiating myoblasts (IFM_d, IFM_e, IFM_f, IFM_g) (Figure S2A, Data S4). The expression of a selected gene panel was visualized by a dot plot (Figure 6I), which allowed us to identify specific markers for each cluster.

### The Argk1 depletion does not alter the identity of the myoblasts in the wing discs

To determine whether the depletion of Argk1 affects the myoblasts across undifferentiated and differentiating clusters, we compared the single cell transcriptomes between *Mef2>Argk1-RNAi* (5,681 myoblasts) and the control *Mef2>mCherry-RNAi* (5,808 myoblasts). In agreement with the *in vivo* results (Figure 1C), the expression of Argk1 was significantly reduced in the Argk1-depleted myoblasts (Figure 6J, Data S5). Distribution of myoblasts of each genotype across cell clusters showed no difference in the contribution of Argk1-depleted cells to each cluster (Figure 6K-L). This was consistent across the two replicate samples (Figure S2B-C). Next, we compared the expression of a known panel of marker genes between Argk1-depleted and control myoblasts across all eight IFM clusters and three DFM clusters (Figure 6M, Data S5). Overall, no significant changes were found in the expression of DFM markers (*kirre* and *ct*) and IFM markers (*zfh1* and *vg*), as well as markers for undifferentiated IFM clusters, (*Him*, *E(spl)m6-BFM*, *E(spl)m3-HLH*, *Sp1*) and for differentiating IFM clusters (*rn*, *h*, *beat-IIb*, *nkd*, *Col4a1*, and *ptc*). We confirmed these results by examining the expression of some of these cluster-specific markers in the adepithelial layer of *Mef2>Argk1-RNAi* wing discs by immunofluorescence. Staining with anti-Ct, anti-Zfh1, anti-H, anti-LamC antibodies and labeling cells with *Him-CD8-GFP*, *Sp1::GFP*, and *E(spl)m6-BFM-gapGFP* reporters revealed no significant changes in the pattern or their expression in Argk1-depleted myoblasts (Figure 6C-G). Thus, the depletion of Argk1 at the onset of flight muscle development does not affect the identity of myoblasts across undifferentiated and differentiating states nor the expression or localization of the well-known myoblast markers.

### Argk1 knockdown affects energy homeostasis in the muscle

To determine the changes in gene expression in the Argk1-depleted myoblasts, we used DE-seq2 to perform a pseudobulk differential gene expression analysis between *Mef2>Argk1-RNAi* and *Mef2>mCherry-RNAi* using the scRNA-seq datasets (Data S5, Data S6). The list of differentially expressed (DE) genes was then subjected to enrichment analyses for gene ontology for biological processes (GOBP), gene ontology for molecular function (GOMF) and KEGG pathways (Data S7). Interestingly, we found that metabolic-related categories were significantly enriched following Argk1 knockdown (Figure 7A). Further, we identified multiple oxidoreductase activity terms in the GOMF enrichment analysis (Figure 7A, middle panel), suggesting possible changes in the NAD+/NADH ratio. To validate this prediction, we measured NAD+/NADH ratio in *Mef2>Argk1-RNAi* and control animals staged at pharate stage using a sensitive fluorescence-based quantification assay. Consistent with the results of the GOMF enrichment analysis, the NAD+/NADH ratio significantly dropped in the Argk1-depleted muscles compared to the control (Figure 7B), thus indicating an increase in the reductive state.

**Figure 7.**
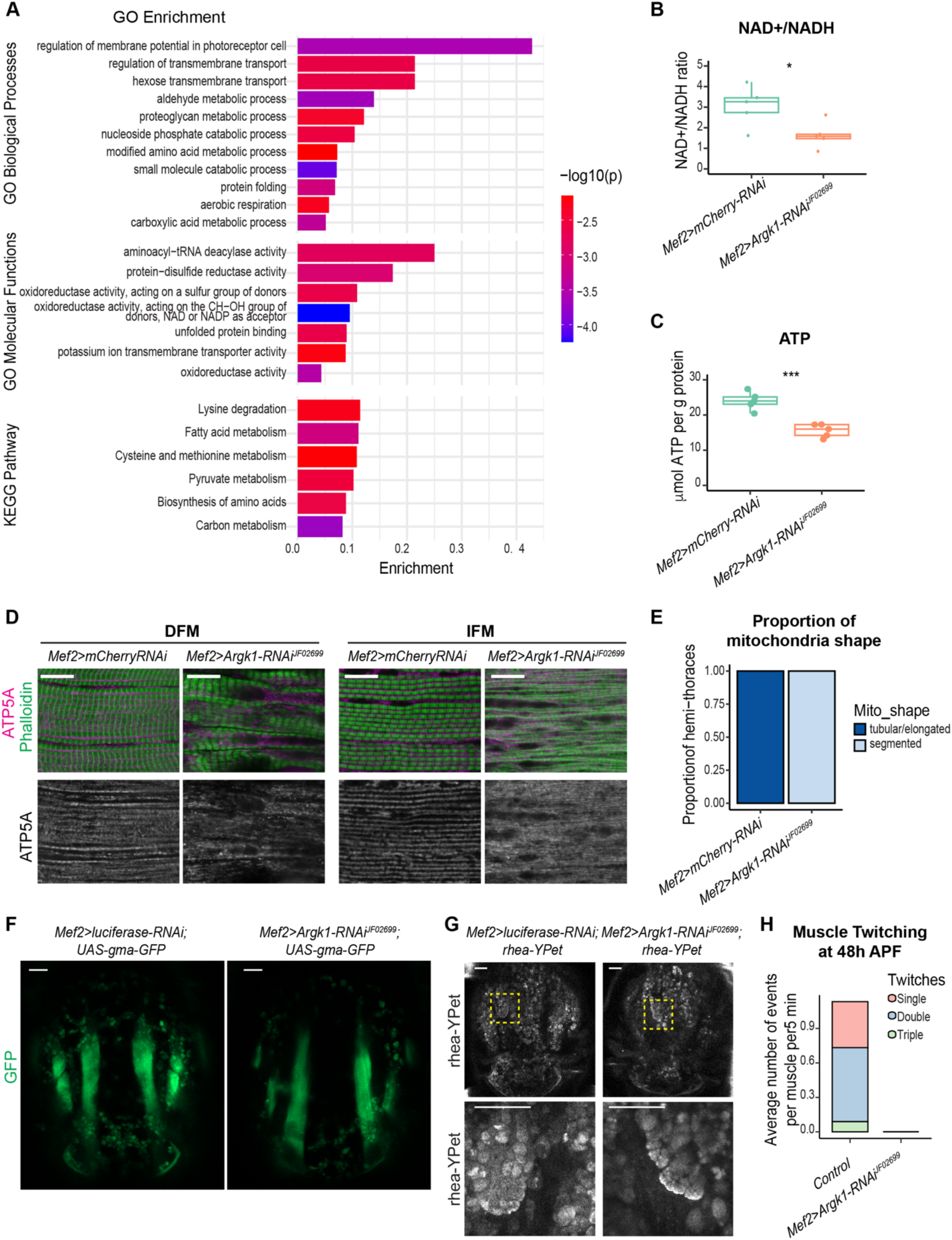
Argk1 is required for the regulation of energy homeostasis in muscles. (**A**) Metascape functional annotation and analysis for the differentially expressed genes between Argk1-depleted myoblasts and control determined by pseudobulk analysis of scRNAseq using DE-seq2. All differentially expressed genes were analyzed together (377 genes, p<0.05). The categories GO term biological processes, molecular processes and KEGG pathway are shown. (**B**) Fluoremetric NAD+/NADH ratio measured on extracts harvested from *Mef2>mCherry-RNAi* and *Mef2-Argk1-RNAi^JF02699^* animals staged at 90h APF (pharates). n = 5 samples/genotype, Welch Two Sample t-test, * p<0.05. N=2 independent experiments. One representative experiment is shown. (**C**) Luminescence-based ATP measured on extracts harvested from *Mef2>mCherry-RNAi* and *Mef2-Argk1-RNAi^JF02699^* animals staged at 90h APF (pharates). Values were normalized to total protein content. n = 5 samples/genotype, Welch Two Sample t-test, *** p<0.001. N=2 independent experiments. One representative experiment is shown. (**D**) Confocal single plane of DFM (muscle 53) and IFM (dorsal longitudinal muscles) examined in *Mef2>Argk1-RNAi^JF02699^* and control *Mef2>mCherry-RNAi* animals staged at 90h APF in a sagittal view. Hemithorax sections stained with Phalloidin (green) and anti-ATP5A. (**E**) Blind quantification of the number of hemi-thoraces displaying either proper morphology of mitochondria (tubular/elongated shape) or segmented mitochondrial shape from IFM in D. Stacked bars. N = 18-20 hemithoraces/genotype. N=2 independent experiments. (**F**) Confocal single plane of live imaging of developing flights muscles from *Mef2>Argk1-RNAi^JF02699^* and control *Mef2>Luciferase-RNAi* pupa staged at 48h APF in a dorsal view. Anterior is to the top. (**G**) Confocal single plane of live imaging of developing flights muscles from *Mef2>Argk1-RNAi^JF02699^* and control *Mef2>Luciferase-RNAi* pupa staged at 48h APF in a dorsal view. Anterior is to the top. Muscle attachment site is labeled with *rhea-YPet*. The yellow-dashed box indicates magnified area for the recorded area in bottom panel. For live movies see Video S1 and Video S2. (**H**) Quantification of spontaneous contraction events per muscle per 5 min. Single, double and triple twitches were counted in developing flight muscles staged at 48h APF from 15 min live recordings from G. Stacked bars. N = 15-16 muscles/genotype. All box plot shows median (middle), interquartile range (box), 1.5x the interquartile range (whiskers), and individual data (points). Scale (**D**) 10 μm, and (**F-G**) 50 μm. Genotypes are: (**A-E**) w; +; *Mef2-GAL4*/*UAS-mCherry-RNAi* w; +; *Mef2-GAL4*/*UAS-Argk1-RNAi^JF02699^* (**F**) w; +; *UAS-Luciferase-RNAi*/*Mef2-GAL4, UAS-gma-GFP* w; +/+; *UAS-Argk1-RNAi^JF02699^*/*Mef2-GAL4,* UAS-*gma-GFP* (**G-H**) w; *Mef2-GAL4/+*; *UAS-Luciferase-RNAi*/*rhea-YPet* w; *Mef2-GAL4/+*; *UAS-Argk1-RNAi^JF02699^*/*rhea-YPet*

We further explored the impact of Argk1 knockdown on energy metabolism by examining the ATP level and mitochondrial morphology in Argk1-depleted muscles. Extracts from animals staged at pharate stage were prepared to measure the total ATP content by a luminescence-based ATP assay, which was then normalized to the total protein content. Notably, the level of ATP was significantly reduced in *Mef2>Argk1-RNAi* animals relative to control (Figure 7C).

To examine mitochondrial morphology, sagittal sections of the DFMs and IFMs were stained with anti-ATP5A antibody to visualize mitochondria and with Phalloidin to label the myofibrils. In the IFMs of control pharates, mitochondria are tightly packed between the myofibrils and form a tubular and elongated mitochondrial network (Figure 7D, right panel). Strikingly, the mitochondrial network was highly abnormal in Argk1-depleted muscles (Figure 7D). The mitochondrial network was segmented and no longer exhibited the tubular shape found in the control. A blind quantification of mitochondrial network shape, categorized as either tubular/elongated or segmented, confirmed that the depletion of Argk1 resulted in statistically significant alterations in mitochondrial morphology (Figure 7E). Similarly, alterations in mitochondria morphology were also found in the DFMs of the Argk1-depleted muscles compared to control ((Figure 7D, left panel). This result was confirmed with another *Argk1-RNAi* transgene *GD10436* (Figure S3A). Consistent with the weaker phenotypes caused by *GD10436* transgene in other tests (Figure 2, Figure S1), the mitochondrial network defects were less severe according to the blind quantification of the mitochondrial shape (Figure S3B).

Overall, these results demonstrate that Argk1 depletion results in severe metabolic abnormalities including changes in the expression of metabolic genes, low ATP level, altered NAD+/NADH ratio and abnormal morphology of the mitochondrial network. Given the interplay and coordination between myofibril morphogenesis and mitochondrial dynamics (*25*), the abnormal shape of the mitochondrial network could be due to defects in the properly forming myofibrils in the Argk1-depleted muscle.

### Argk1-depleted muscles fail to undergo spontaneous contractions during pupal development

Muscles undergo spontaneous contractions during development, which is important for muscle formation (*26*). The contractions can be first seen at the beginning of the sarcomere assembling in immature myofibrils at 30h APF, with the peak at 48h APF, and at 72h APF the contractions cease (*21*). To determine whether Argk1 knockdown affects these spontaneous contractions, we performed live imaging microscopy of IFM in intact pupa at 48h APF carrying transgenes: *Mef2>gma-GFP* to label muscles (Figure 7F) and *rhea-YPet* to label the muscle attachment sites for visualization of the twitching events (Figure 7G, Video S1). In control pupa, about one twitching event consisting of either single, double or triple contractions was recorded every 5 minutes (Figure 7H). Strikingly, spontaneous contractions were completely absent in Argk1-depleted muscles. We did not detect a single twitching event in *Mef2>Argk1-RNAi* animals (Figure 7H, Video S2). Significantly, the animals were alive at that time point since very mild movements across all flight muscles were readily observed occasionally.

Thus, Argk1-depleted muscles were unable to undergo spontaneous muscle contraction during pupal development, which are important for sarcomere assembly and myofibrillogenesis. Given that the ATP level is low in these animals, the lack of muscle twitching suggests that Argk1 is required in the muscle to buffer ATP levels and regulate energy homeostasis during muscle formation.

## Discussion

The analysis of the transcriptional profiles for the wing disc associated myoblasts by scRNA-seq offers a way to identify novel genes that are functionally important for early stages of flight muscle development. Indeed, many of the scRNA-seq derived markers for individual myoblast clusters were also isolated in a large-scale muscle-related RNAi screen (*18*). In this work, we focused on *Argk1* that was highly expressed across several myoblast clusters and caused animal lethality upon its knockdown in the muscle (*9*). Argk1 is well known for its importance in ATP buffering in cells with highly volatile power demands, such as contracting muscles (*27*). Here, by examining the developing Argk1*-*depleted muscles, we uncovered a novel role for Argk1 in maintaining energy homeostasis during flight muscle development that is required for muscle growth and myofibrillogenesis.

Argk1 belongs to the group of phosphagen kinases responsible for catalyzing the reversible transfer of a phosphate group between ATP and arginine, resulting in the formation of phosphorylarginine. Since ATP is readily catalyzed by the phosphagen kinase during high energy turnover (*28*, *29*), phosphagens, such as phosphorarginine, serve as an on-demand reservoir of “high-energy phosphates”. The phosphagen kinases are important in proton buffering and in intracellular energy transport and play a dynamics role in the metabolic regulation of cells with high and variables rates of energy turnover, including muscles, neurons and spermatozoa (*30*). In invertebrates, the activity of Arginine kinase has been associated with ATP regeneration and energy transport during burst of ATP demand in tissues with high level of energy metabolism (*27*, *30*, *31*). The flight muscles is one of the highest metabolically demanding organs, as they undergo a 50-100-fold increase in oxygen consumption from rest to flight. Contracting flight muscles rely mainly on aerobic glycolysis and oxidative phosphorylation to generate ATP (*32*, *33*). To maximize the efficiency of ATP delivery to the myosin ATPase for muscle contraction, the glycolytic enzymes are located within the sarcomere of the myofibrils (*34*, *35*). Similarly, Argk1 is highly abundant in skeletal muscles (*12*, *14*) and is present within the sarcomere structures of the flight muscles that ensures localized ATP availability for muscle contraction (*13*). Thus, Argk1 is commonly viewed as an important player in maintaining energy homeostasis during periods of high demand or burst of energy fluctuation to buffer the ATP levels. Our data are consistent with this established role of Argk1 since its depletion late in development leads to impaired muscle function in adults, which can be seen by the reduced performance of these animals in the flight test assay and by mild defects in sarcomere assembly and myofibrillogenesis. However, this view of Argk1 function appears to be incomplete, as our findings reveal that Argk1 is also required throughout muscle development. This conclusion is supported by the reduced muscle size and severe defects in myofibrillogenesis in both direct and indirect flight muscles when Argk1 is depleted using early muscle specific GAL4 drivers. Accordingly, immunofluorescence and scRNA-seq show that Argk1 is highly expressed throughout flight muscle development, and therefore its expression is not limited to adults. Despite severe muscle defects, Argk1-depleted myoblasts turn on the proper transcriptional programs as they initiate differentiation at earlier stages of muscle development. We also did not find any alterations in cell cycle progression or cell division of proliferating myoblasts when Argk1 is depleted. However, Argk1 knockdown results in abnormal NAD+/NADH ratio and strong reduction of ATP levels, which is consistent with the role of Argk1 in maintaining energy homeostasis. Accordingly, Argk1-depleted myoblasts are smaller in size. Thus, our data support the idea that Argk1 is needed to buffer ATP levels during flight muscle development.

Recent works characterized spontaneous contractions of the flight and abdominal muscles during their development (*21*, *26*). Such muscle twitching events are important since changes in the frequency of contractions affects the myofibrillogenesis. Interestingly, we found that Argk1-depleted muscles completely lack such twitching events and, concordantly, have abnormal sarcomere structures. This finding suggests that Argk1 is required for spontaneous muscle twitching during development presumably by supporting high energy demand during contractions, a function that is highly reminiscent to the established role of phosphagen kinases in adult muscle. As spontaneous muscle twitching is a required event for myofibrillogenesis (Spletter 2018, and their absence may explain severe defects in myofibrillogenesis and sarcomere assembly in Argk1-depleted muscle.

Mitochondrial morphogenesis is tightly coordinated with myofibril assembly and is mediated by mechanical tensions and kinesin-dependent microtubules (*25*, *36*), while mitochondrial fission and fusion are spatiotemporally regulated during muscle differentiation and growth (*37*). Interestingly, the mitochondrial network is highly fragmented in Argk1-depleted muscle. Given that myofibrils mechanically constrain mitochondria into elongated shape in flight muscle (*25*) the defects in mitochondrial morphology observed in Argk1-depleted muscle could be the consequence of disrupted myofibril assembly. Overall, our data is in concordance with previous findings that highlight the importance in coordinating myofibril assembly with mitochondria morphogenesis.

In summary, our data suggest that Argk1 is important during muscle development to maintain the ATP level, which is critical for spontaneous muscle twitching and other processes with high energy demand during myofibrillogenesis and muscle growth. Thus, *Argk1* plays a broad role in energy homeostasis that impacts both normal muscle physiology and proper muscle formation.

## Data availability

Chromium10x scRNA-seq data have been deposited to the NCBI Gene Expression Omnibus database and are accessible through the accession number GSE138625 (https://www.ncbi.nlm.nih.gov/geo/query/acc.cgi?acc=GSE138625).

## Acknowledgments

We thank Holly Jefferson for technical help, Richard Cripps for sharing fly stock Act88F-GAL4, Maria Spletter and Frank Schnorrer for sharing the rhea-YPet and Him-CD8-GFP stocks, and K. Jagla for E(spl)m6-BFM-gap-GFP. We thank Richard Cripps and Maria Spletter for discussion. Other stocks were obtained from the Bloomington Drosophila Stock Center (NIH P40OD018537), the Vienna Drosophila Resource Center, the TRiP at Harvard Medical School. We are grateful to Developmental Studies Hybridoma Bank and the Babraham Institute for antibodies, Flybase for online resources on the Database of Drosophila Genes and Genomes. We thank the Single Cell Sequencing Pilot at University of Illinois at Chicago and DNA services Core at University of Illinois at Urbana-Champaign for sequencing. This work was supported by NIH grant R35GM131707 to M.V.F.

## Author contributions

M.P.Z. and M.V.F. conceived the project, designed the experiments, analyzed data, and wrote the manuscript. M.P.Z., A.W., H.C., R.G., L.T., L.dC., and O.C. performed and generated the data. All authors reviewed the manuscript.

