## Supplemental figures and data for "*Arginine Kinase 1* regulates energy homeostasis in *Drosophila* muscle development"

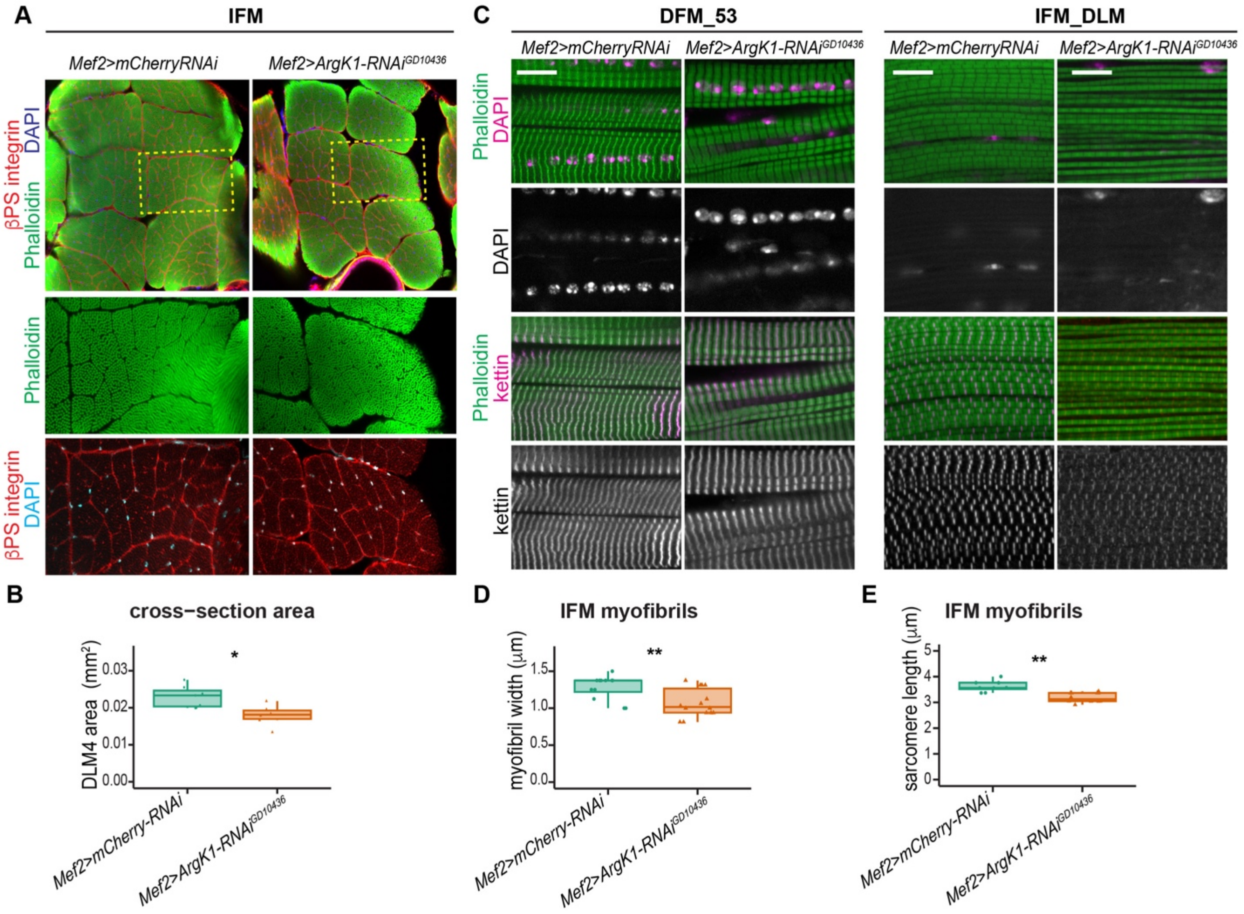


**Figure S1. Argk1 knockdown driven by *Mef2-GAL4* results in severe defects in both IFM and DFM.** Related to Figure 3

(A-E) The structure of the DFM and IFM examined in *Mef2>Argk1-RNAi*$^{GD10436}$ and control *Mef2>mCherry-RNAi* animals at 1 or 2-days-old. (A) Confocal single plane images of IFM in a transverse view of the dorsal longitudinal muscles (DLM). Sections stained with Phalloidin (green), DAPI (blue) and βPS-integrin (red). Yellow-dashed box indicates magnified area for DLM4 (bottom panel). Dorsal up. (B) Quantification of the cross-sectional area for the DLM4 per animal. N = 6-7 thoraces/genotype, Wilcoxon rank-sum test, * $p<0.05$. (C) Confocal single plane images of DFM (Muscle 53, left panel) and IFM (dorsal longitudinal muscles, right panel) in a sagittal view. Hemithorax sections stained with Phalloidin, DAPI and anti-Kettin. (D) Quantification of myofibril diameter for the IFM-DLM per animal. N = 12-14 thoraces/genotype, Wilcoxon rank-sum test, ** p<0.01. (E) Quantification of sarcomere length for the IFM-DLM per animal. N = 8-12 thoraces/genotype, Wilcoxon rank-sum test, ** p<0.01.

All box plot shows median (middle), interquartile range (box), 1.5x the interquartile range (whiskers), and individual data (points).

Scale (A) 50 μm, and (C) 10 μm.

Genotypes are:

w; +; *Mef2-GAL4/UAS-mCherry-RNAi*

w; +; *Mef2-GAL4/UAS-Argk1-RNAi*$^{GD10436}$

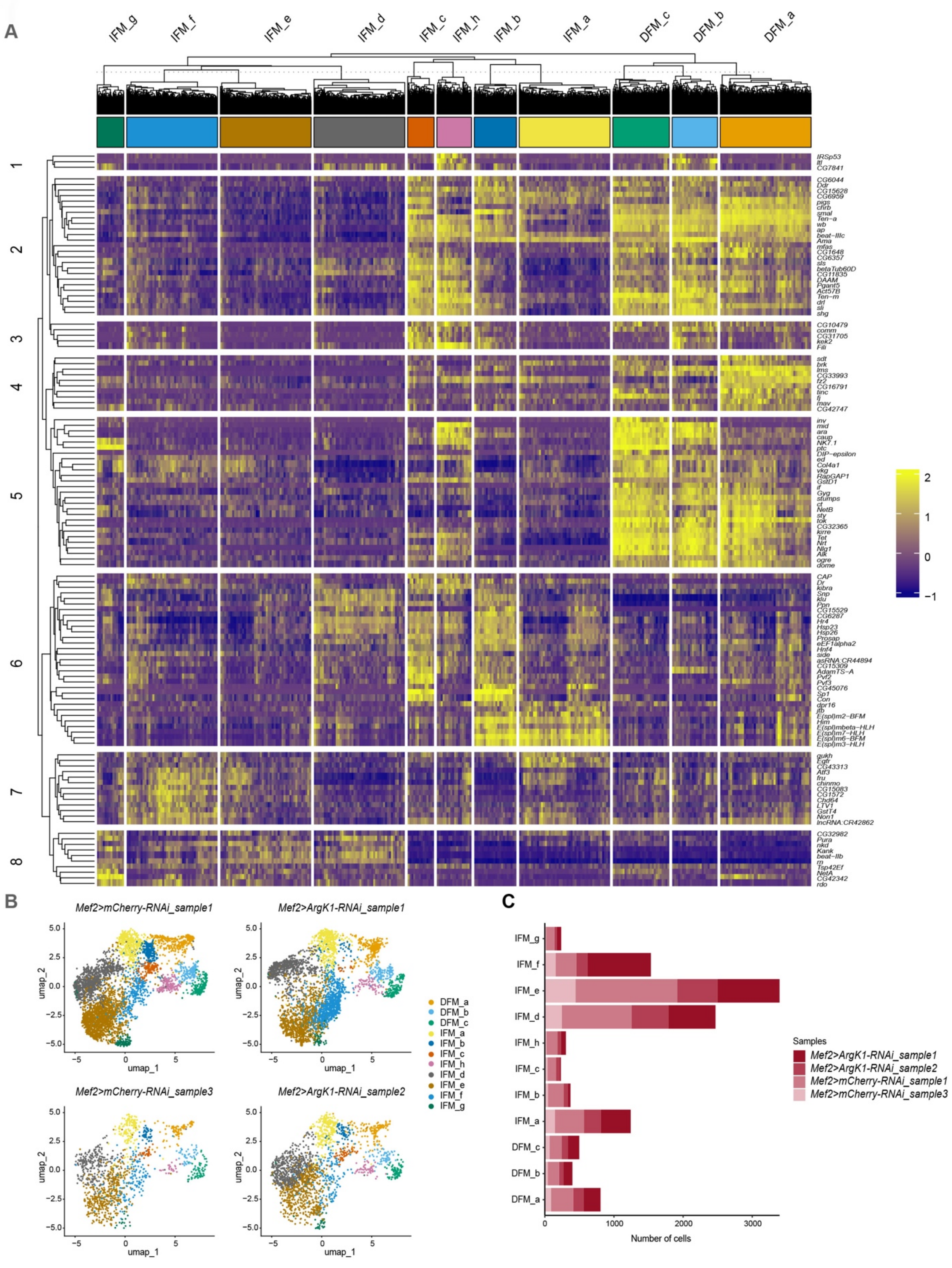
A
IFM_g
IFM_f
IFM_e
IFM_d
IFM_c
IFM_h
IFM_b
IFM_a
DFM_c
DFM_b
DFM_a
1
2
3
4
5
6
7
8
IRSp53
ftl
CG7841
CG6044
Ddr
CG15628
CG6959
pigs
chrb
smal
Ten-a
wb
ap
beat-IIIc
Ama
mfas
CG1648
CG6357
sls
betaTub60D
CG11835
DAAM
Pgant5
Act57B
Ten-m
drl
sli
shg
CG10479
comm
CG31705
kek2
Fili
sdt
brk
lms
CG33993
fz2
CG16791
tinc
fj
mav
CG42747
inv
mid
ara
caup
NK7.1
ptc
DIP-epsilon
ed
Col4a1
vkg
RapGAP1
GstD1
if
Gyg
stumps
ct
NetB
sty
tok
CG32365
kirre
Tet
Nrt
Nlg1
Alk
ogre
dome
CAP
Dr
kibra
Snp
klu
Ppn
CG15529
CG6287
Hr4
Hsp23
Hsp26
Prosap
eEF1alpha2
Hnf4
side
asRNA:CR44894
CG15309
AdamTS-A
Pvf2
Pvf3
CG45076
Sp1
Con
dpr16
ftb
E(spl)m2-BFM
Him
E(spl)mbeta-HLH
E(spl)m7-HLH
E(spl)m6-BFM
E(spl)m3-HLH
gukh
Egfr
CG43313
Atf3
fru
chinmo
CG15083
CG1572
Chd64
LTV1
GstT4
Non1
lncRNA:CR42862
CG32982
Pura
nkd
Kank
beat-IIb
rn
Tsp42Ef
NetA
CG42342
rdo
2
1
0
−1
B
Mef2>mCherry-RNAi_sample1
Mef2>ArgK1-RNAi_sample1
Mef2>mCherry-RNAi_sample3
Mef2>ArgK1-RNAi_sample2
umap_1
umap_2
DFM_a
DFM_b
DFM_c
IFM_a
IFM_b
IFM_c
IFM_h
IFM_d
IFM_e
IFM_f
IFM_g
C
Number of cells
0
1000
2000
3000
Samples
Mef2>ArgK1-RNAi_sample1
Mef2>ArgK1-RNAi_sample2
Mef2>mCherry-RNAi_sample1
Mef2>mCherry-RNAi_sample3

**Figure S2. Argk1 knockdown does not alter the overall identity of the undifferentiated and differentiated myoblasts.** Related to Figure 6

(A) Hierarchical heatmap showing the expression levels of 124 genes listed as the top 30 markers per cluster (adjp<0.05, log2FC>1, pct1>0.5). Hierarchical clustering by k-means to group both the clusters of cells and the genes into groups based on similarity. Color intensity shows the average expression level normalized and scaled.

(B) Two-dimensional UMAP representation of both the combined *Mef2>Argk1*- RNAi$^{JF02699}$ (5,681 myoblasts) and *Mef2>mCherry-RNAi* (5,808 myoblasts) from third instar larval wing discs datasets, colored by cell cluster and split by replicate.

(C) Total number of cells per cluster per sample relative to the total number of myoblasts per sample for the combined datasets *Mef2>Argk1*- RNAi$^{JF02699}$ and *Mef2>mCherry-RNAi* across four samples.

Genotypes are:

w; +; *Mef2-GAL4/UAS-mCherry-RNAi*

w; +; *Mef2-GAL4/UAS-Argk1-RNAi*$^{JF02699}$

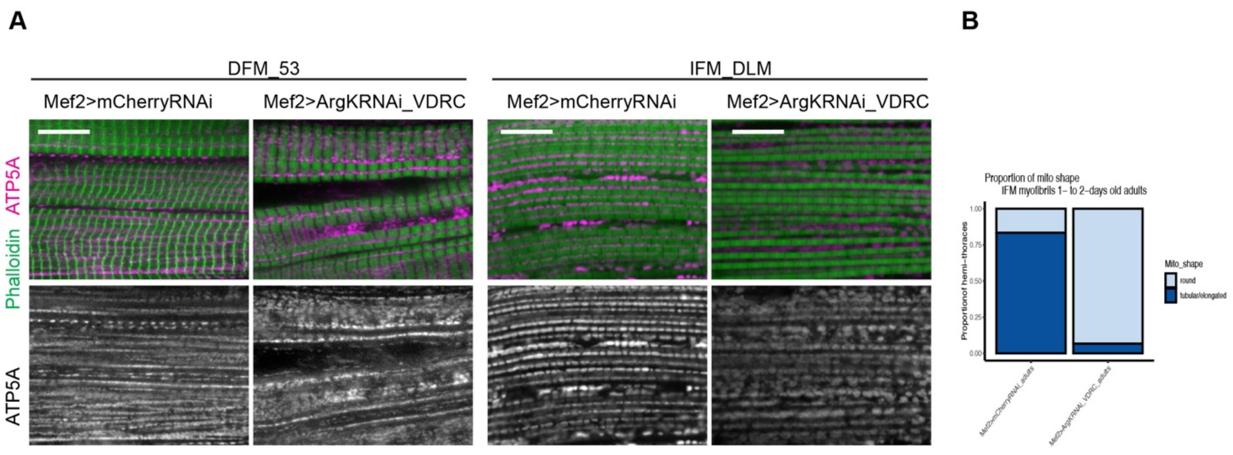


**Figure S3. Argk1 is required for the regulation of energy homeostasis in muscles.** Related to Figure 7.

(**A**) Confocal single plane of DFM (muscle 53, left panel) and IFM (dorsal longitudinal muscles, right panel) examined in *Mef2>Argk1-RNAi*$^{GD10436}$ and control *Mef2>mCherry-RNAi* animals staged at 1 or 2-days old in a sagittal view. Hemithorax sections stained with Phalloidin (green) and anti-ATP5A. Scale 10 μm.

(**B**) Blind quantification of the number of hemi-thoraces displaying either proper morphology of mitochondria (tubular/elongated shape) or round/segmented mitochondrial shape from IFM in A. Stacked bars. N = 15-18 hemithoraces/genotype. N=2 independent experiments.

Genotypes are:

w; +; *Mef2-GAL4/UAS-mCherry-RNAi*

w; +; *Mef2-GAL4/UAS-Argk1-RNAi*$^{GD10436}$

**Movie S1**. **Live imaging of *Mef2>Luciferase-RNAi* developing flights muscles**. Related to Figure 7.

Animal staged at 48h APF recorded in a dorsal view. Anterior to top. Muscle attachment site labeled with rhea-YPet. Movie created at 20 frames per second.

Genotype is *w; Mef2-GAL4/+; UAS-Luciferase-RNAi/rhea-YPet*

**Movie S2**. **Live imaging of *Mef2>Argk1-RNAi*$^{JF02699}$ developing flights muscles.** Related to Figure 7.

Animal staged at 48h APF recorded in a dorsal view. Anterior to top. Muscle attachment site labeled with rhea-YPet. Movie created at 20 frames per second.

Genotype is *w; Mef2-GAL4/+; rhea-YPet/UAS-Argk1-RNAi*$^{JF02699}$

**Data S1. List of cell barcodes used after applying quality control filters and selecting the myoblasts**. Related to Figure 6.

**Data S2. Markers for each cluster of the combined dataset *Mef2>mCherry-RNAi* and *Mef2>Argk-RNAi* myoblasts**. Related to Figure 6.

List of all differentially expressed genes across all clusters. Only positive markers with threshold set p_val_adj<0.05

**Data S3. Average expression of all genes across all clusters of the combined dataset *Mef2>mCherry-RNAi* and *Mef2>Argk-RNAi* myoblasts**. Related to Figure 6.

List of average gene expression across all clusters (SCT assay).

**Data S4. Top biomarkers per cluster grouped into 8 groups by k-means of the combined dataset *Mef2>mCherry-RNAi* and *Mef2>Argk-RNAi* myoblasts**. Related to Figure 6.

The top unique 30 markers per cluster (124 genes, adjp<0.05, log2FC>1, pct1>0.5) clustered by k-means to group genes based on similarity.

**Data S5. Aggregate Expression across Genotypes and across Samples of the combined dataset *Mef2>mCherry-RNAi* and *Mef2>Argk-RNAi* myoblasts**. Related to Figure 6.

Aggregate Expression across the two genotypes and across all four samples

**Data S6. Differential expression using pseudobulk (DESeq2) between *Mef2>mCherry-RNAi* and *Mef2>Argk-RNAi* myoblasts**. Related to Figure 7.

List of differentially expressed genes between *Mef2>mCherry-RNAi* and *Mef2>Argk1-RNAi* (DE-Seq2, p<0.05, 377 genes). Genes are listed by their gene number in FlyBase (FBgn).

**Data S7. Metascape functional annotation and analysis.** Related to Figure 7.

Functional annotation and enrichment analysis by Metascape for the differentially expressed genes between *Mef2>mCherry-RNAi* and *Mef2>Argk-RNAi* myoblasts determined by pseudobulk analysis (DE-seq2, 377 genes, p<0.05). The categories GO term biological processes, molecular processes and KEGG pathway are listed.
